# Demand for Multiplatform and Meta-analytic Approaches in Transcriptome Profiling

**DOI:** 10.1101/860312

**Authors:** Dóra Tombácz, Gábor Torma, Gábor Gulyás, Norbert Moldován, Michael Snyder, Zsolt Boldogkői

## Abstract

In a recent article, Depledge and colleagues reported a study of the herpes simplex virus type 1 (HSV-1) transcriptome using direct RNA sequencing (dRNA-Seq) on nanopore arrays. The authors provided a useful dataset on full-length viral and host RNA molecules. In this study, we reanalyzed the published dataset and compared it with data generated by our group and others. Our comparative study clearly demonstrated the need for multiplatform and meta-analytic approaches for transcriptome profiling to obtain reliable results.

## Introduction

Second-generation short-read sequencing (SRS) technology, launched in the mid-2000s, has revolutionized both genomic and transcriptomic researches because of its ability to sequence millions of nucleic acid fragments simultaneously at a relatively low expenditure per base. In recent years, the third-generation long-read sequencing (LRS) approaches have been emerged. Currently, two LRS approaches are in use: the single-molecule real-time technology developed by Pacific Biosciences (PacBio) and the nanopore-based sequencing developed by Oxford Nanopore Technologies (ONT). LRS can overcome several shortcomings of SRS in transcriptome analysis, which is mainly based on the ability of these techniques to read full-length RNA molecules. However, similarly to SRS, LRS techniques often produce spurious transcripts owing to issues such as template switching and mispriming in reverse transcription (RT) and PCR. The major problem is that no efficient bioinformatic tools are currently available to detect these errors. Native RNA sequencing has been considered superior to cDNA sequencing because of the lack of artifacts generated via amplification of RT and PCR (however, notably, direct cDNA sequencing without PCR amplification is also possible using both LRS platforms). Nonetheless, dRNA-Seq has also limitations, such as low throughput, a lack of 15-30 bases from the transcription start site, and errors produced, for example by the ligation used for the attachment of adapters, by the single-strand cDNA formation, or by the potential slippage of RNA molecules during their passage across the nanopore as a result of temporary improper functioning of the ratcheting enzyme. The low throughput of dRNA-Seq makes both the transcript identification and the annotation of nucleic acid sequences at base-pair resolution difficult, which is especially critical in large-genome species.

LRS has already been applied for the transcriptome analysis of various organisms ^1,2^, including herpesviruses ^3–6^. This approach has revealed extremely complex transcriptome profiles in every examined species. LRS techniques can be used in analyses that are challenging for SRS approaches, such as the detection of multi-spliced transcripts, parallel transcriptional overlaps, low-abundance transcripts, and very long and embedded RNA molecules. A single technique may fail to detect certain transcripts or transcript isoforms, and to precisely map the transcript ends or the intron boundaries. Additionally, the platform- and library preparation-dependent sequencing errors may produce false isoforms. A meta-analysis including multiplatform approaches, such as various LRS and SRS techniques, as well as different auxiliary methods, such as cap selection, and 5’- and 3’-ends mapping, can circumvent this problem, especially if different library preparation protocols are used. Furthermore, the comparison of the various data provides a tool for identifying novel transcripts, validating already-described RNA molecules, or removing putative transcripts if not confirmed by other techniques.

## Results

In this study, we employed an integrated approach based on the meta-analysis of the HSV-1 transcriptome data published by Depledge and colleagues (using ONT dRNA-Seq and Illumina RNA-Seq)^7^, Tang et al. (using Illumina SRS)^8^, Rutkowski et al. (using Illumina SRS)^9^, Wishnant et al. (using Illumina SRS)^10^, and our laboratory (Tombácz and colleagues using PacBio RSII^11^, as well as Boldogkői et al.^12^ and Tombácz et al.^13^ using PacBio Sequel, ONT dRNA-Seq and cDNA sequencing with multiple library preparation methods; **Supplementary Table 1**). This analysis led to the discovery of novel transcripts, especially of novel multigenic transcripts (**Supplementary Figure 1**), and splice sites (**Figure 1, Supplementary Figure 2)**. As Figure 1 shows, a relatively high percentage of introns were not detected in other studies, for which the probable reason is the extremely strict criteria for the annotations. Additionally, we confirmed putative RNA molecules and transcripts isoforms, which were previously unpublished because of inadequate evidence supporting their existence (**Supplementary Table 2**). This analysis also revealed that practically all HSV-1 genes contain at least one shorter transcript variant with truncated in-frame ORFs (**Figure 2)**. Loosening the annotation criteria probably would lead to the identification of truncated genes in every canonical gene. We also identified several fusion genes with relatively long introns spanning the gene boundaries. Additionally, a large number of low-abundance transcript isoforms, including splice and length variants were identified in this and also in other studies^14^. Whether these molecules have functional significance, or they are merely the result of transcriptional noise remains to be ascertained. The general functions of the embedded and the fusion genes are also unknown. We demonstrated that dRNA sequencing produces a certain level of errors, because, for example, we could not detect a large number of dRNA introns (299 introns in Depledge’s dataset and a single intron in our dRNA-Seq dataset) in either cDNA database, which might be explained by the differences in the coverages. However, the most abundant introns were present in both databases. This study also revealed that using different reference genomes for mapping the same transcripts can lead to somewhat different results with respect to the splice sites, especially in SRS.

**FIGURE 1.**
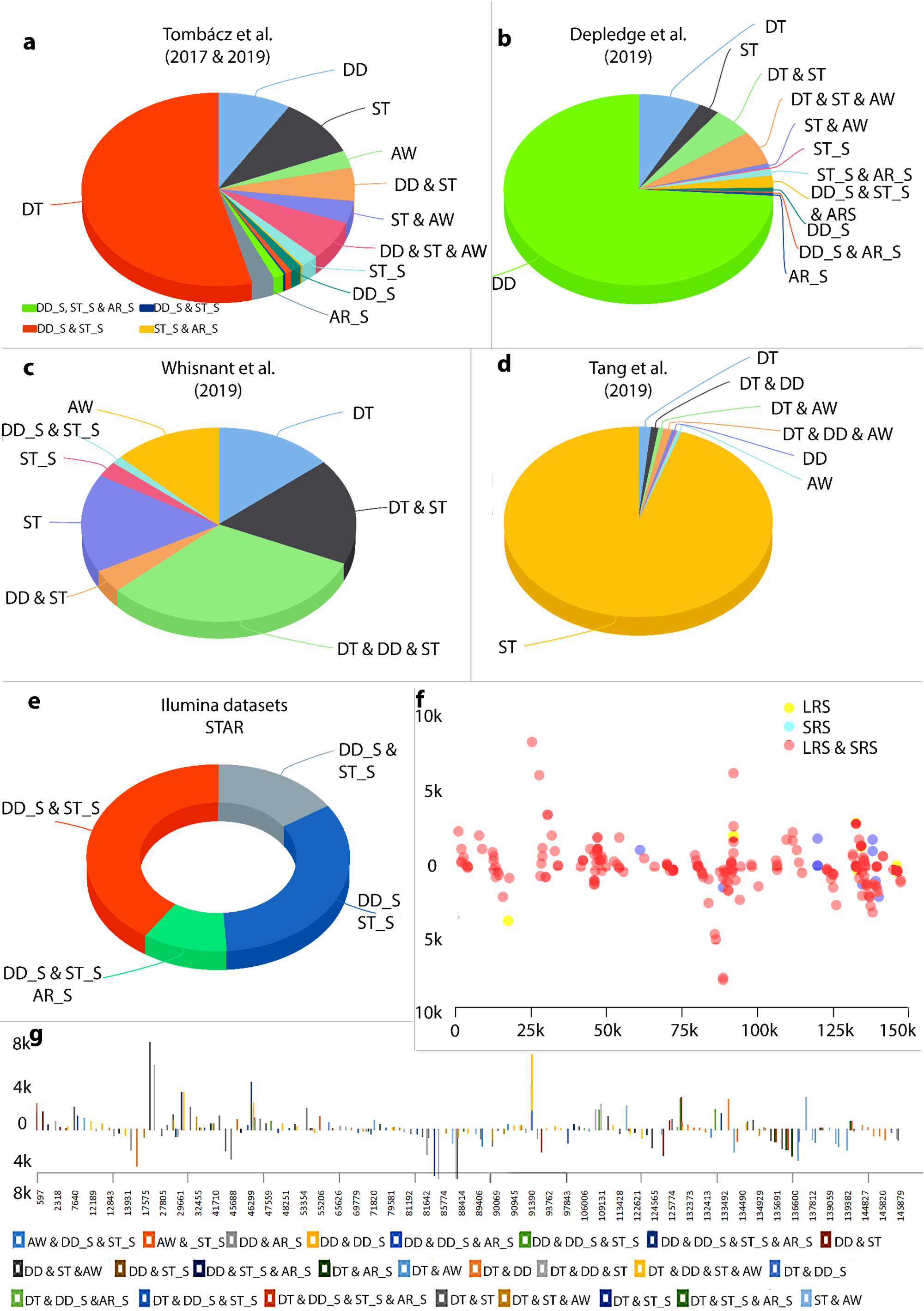
Herpes simplex virus type 1 (HSV-1) introns identified using different sequencing platforms. The 378 putative introns identified in our earlier study^11,13^ are already multiplatform-based (various combination of library preparation techniques of Pacific Biosciences RSII and Sequel, and Oxford Nanopore Technologies MinION sequencing). These datasets were compared with the intron datasets generated by Tang et al.^8^ and Whisnant et al.^10^. We also used the raw sequencing reads from Depledge’s direct RNA-Seq study. The data were aligned to the HSV-1 genome and then analyzed using LoRTIA. This analysis detected 214 introns. Three large raw Illumina datasets^7–9^ were also mapped and reanalyzed. Only the introns that were present in at least two independent datasets were accepted and plotted. We obtained 975 additional introns from this part of the work. **a. Introns identified by Tombácz and colleagues**. Altogether, 45.5% of these introns have been validated by the other studies. **b. Introns identified in Depledge and coworkers’ dataset using the LoRTIA tool.** Our analysis of the raw dRNA-Seq reads detected 437 potential introns, from which 114 were also found in the other studies. The LoRTIA tool did not identify the previously published intron within the RNA encoding the fusion protein RL2–UL1^7^; however, it was verified by Tang and colleagues^8^. **c. Introns from Whisnant and colleagues’ publication.** They have been published 79 introns, 87% of which were also found in the other datasets. The authors have analyzed our previous dataset^11^ and found that seven of the eleven published introns are low-abundance isoforms. Therefore, they considered them as unconfirmed. We found and validated five out of these seven introns in our novel dataset, and they were also present in either Tang’s and/or Depledge’s dataset. **d. Introns published by Tang and colleagues.** These authors published a large number of introns (2352), but only 5% of them were validated in the other datasets. **e. Reanalysis of HSV datasets from various Illumina sequencing experiments.** This work yielded 975 introns, which were detected in at least two of the datasets. **f. Intron lengths.** This scatter plot represents the genomic locations and lengths of the above 214 introns. **g. Intron length.** The colored bar charts show the location and lengths of the introns. The colors represent the various combinations of the techniques by which the given intron was detected. Abbreviations: DT: Tombácz et al. 2017 & 2019; DD: Depledge et al. 2019; ST: Tang et al. 2019; AW: Whisnant et al. 2019; AR_S: dataset from Rutkowski et al. 2019 analyzed by STAR; DD_S: Illumina dataset from Depledge et al. 2019 analyzed by STAR; ST_S: dataset from Tang et al. 2019 analyzed by STAR

**FIGURE 2.**
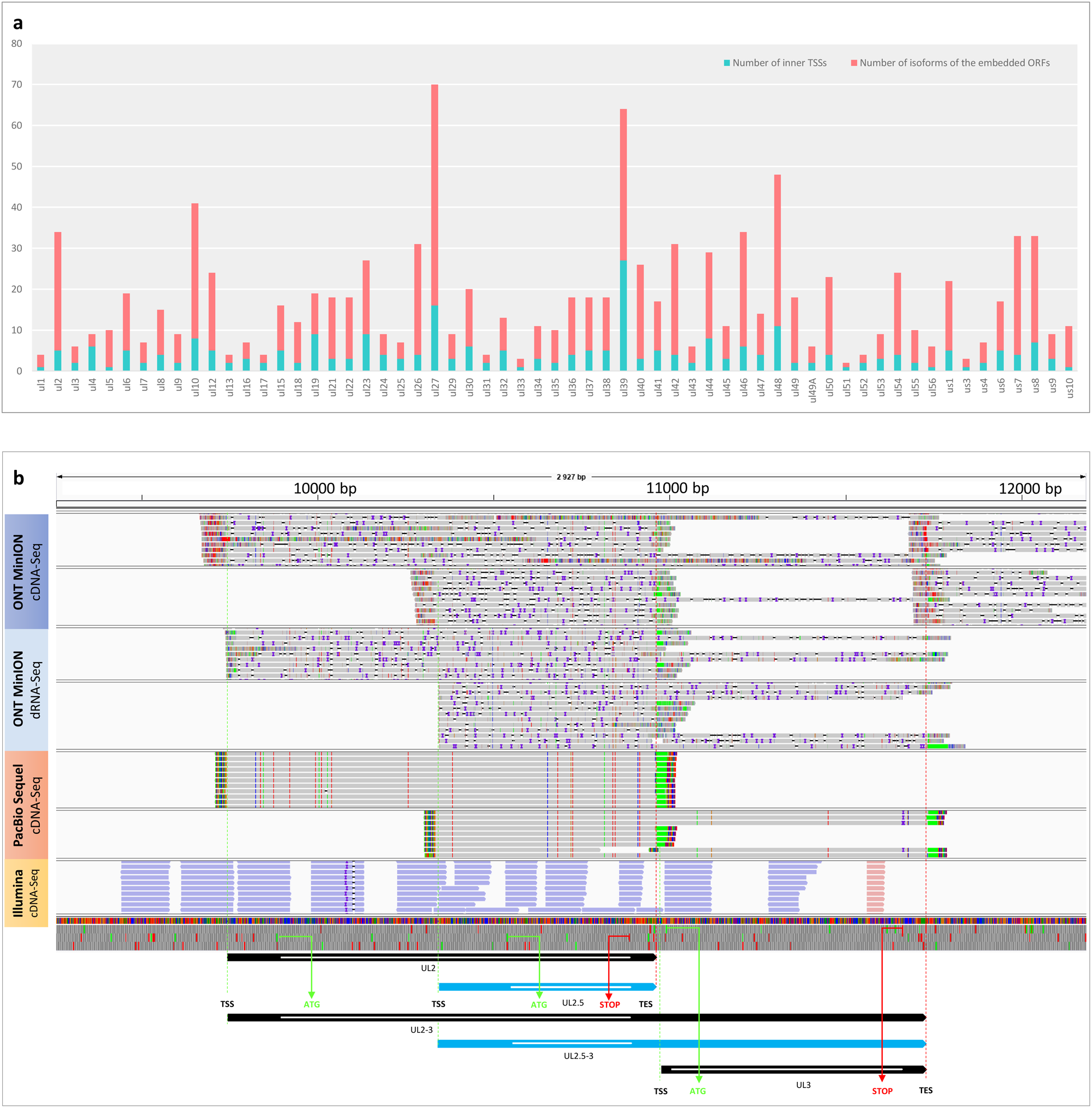
We have published the general occurrence of embedded genes (63 genes) in the herpes simplex virus type 1 genome. Sixty-one of them were validated by the dataset from Depledge’s publication. **a. Bar chart representation of the embedded ORFs.** Many of the embedded ORFs have multiple isoform length, however, this phenomenon is presented in **Supplementary Table 2. b. An example for an embedded ORFs-containing transcript detected by various techniques.** Visualization of the UL2 transcript and one of its truncated transcripts (ul2.5) using Integrative Genomics Viewer. The sequencing reads are from long-read (LRS) sequencing and short-read sequencing (SRS) datasets including direct RNA (dRNA) and cDNA sequencing. It can be seen that the dRNA-seq and the two LRS cDNA techniques detected the same TSS (note that dRNA sequencing produces shorter 5’-UTRs [on average, 23□bp are missing]). The figure also shows that the SRS without a specialized library preparation method (e.g., CAGE) is not sufficient to identify 5’-ends of transcripts.

## Discussion

Taken together, employing multiplatform approaches with distinct library preparation methods is especially important in transcriptome research because of the high error-rate and the variances in the results obtained using miscellaneous library preparation, sequencing and annotation methods. Furthermore, meta-analyses can control the potential errors derived from using different kits and protocols, as well as from dissimilar working styles and conditions in different laboratories.

## Methods

### Datasets

The datasets generated by Depledge et al.^7^ and five other datasets (Tombácz et al.^11,13^; Tang et al.^8^; Rutkowski et al.^9^, and Whisnant et al.^10^) were reanalyzed in order to define the complete HSV transcriptome.

### Data analysis

The adapter sequences from the raw reads of each SRS run were removed by using Cutadapt v2.6 software. The fastp tool was used for validation. Further, we aligned the sequencing reads to the HSV-1 reference genome (GenBank: X14112.1) using minimap2 or STAR mapper for the LRS or the SRS data, respectively. The LoRTIA tool was used to annotate introns and TSSs, and TESs from the LRS data, whereas we used the STAR software was used to detect introns from the SRS samples. The previously published introns (Tang et al.^8^, Wishnant et al.^10^, and Tombácz et al.^11,13^) were compared with each other, reanalyzed, and validated by using the datasets from all of the aforementioned publications.

## Data availability

The datasets used in this work were obtained from the original publications: Depledge et al.,^7^, Whisnant et al.^10^, Tang et al.^8^, Rutkowski et al.^9^, and from Tombácz et al.^11,13^. All data generated in this study are included in **Supplementary Table 1**. The data of introns plotted in this study were obtained from Tang et al.^8^, Rutkowski et al.^9^, and from Tombácz et al.^11,13^. The codes for the LoRTIA (the toolkit developed by our laboratory) analysis are available at: https://github.com/zsolt-balazs/LoRTIA.

## Acknowledgements

We would like to thank Marianna Ábrahám (University of Szeged) for her technical assistance. This study was supported by grants from the National Research, Development and Innovation Office OTKA K 128247 to ZBo and National Research, Development and Innovation Office (OTKA FK 128252 to DT).

## Author contributions

D.T. and Z. B. conceived the idea. D.T., G.T., G.G., N.M., and Z.B. conducted the analysis. D.T., M.S., and Z.B. designed the methodology. D.T. and G.T. prepared the Figures. Z.B. and D.T. wrote the manuscript with feedback from all co-authors. Z.B. and M.S. coordinated the project.

## Additional information

### Conflict of interest

The authors declare no conflict of interest.

## SUPPLEMENTARY MATERIALS

**Global herpes simplex type 1 transcriptome assembled by meta-analysis of different sequencing approaches**

**Supplementary Table 1. Summary table of the sequencing reads aligned to the herpes simplex virus type 1 reference genome. a.** Data of the read count and average read length from the long-read sequencing techniques, using the LoRTIA tool. **b.** Total read count and read length from the short-read sequencing, based on our reanalysis.

**Supplementary Table 2. Herpes simplex virus type 1 (HSV-1) transcripts and introns. a.** Updated transcript list of the HSV-1 virus without the spliced transcripts. **b.** Updated intron list. Abbreviations: DT: Tombácz et al. 2017 & 2019; DD: Depledge et al. 2019; ST: Tang et al. 2019; AW: Whisnant et al. 2019; AR_S: dataset from Rutkowski et al. 2019 analyzed by STAR; DD_S: Illumina dataset from Depledge et al. 2019 analyzed by STAR; ST_S: dataset from Tang et al. 2019 analyzed by STAR. **c**. List of super-long transcripts.

**Supplementary Figure 1. Super-long transcripts of herpes simplex virus type 1.** These large (≥ 4 kbps) RNA molecules were identified by ONT MinION dRNA-Seq and PacBio Sequel techniques. Many of them rare with uncertain TSSs especially those ones which were detected by dRNA-Seq. Only the longest transcripts are illustrated at a certain genomic region except the overlapping transcripts are complementary to each other.

**Supplementary Figure 2. Integrative Genomics Viewer representation of the intron positions.**

