## Supplementary figures and images for "Demand for Multiplatform and Meta-analytic Approaches in Transcriptome Profiling"

### Supplementary Figure 1

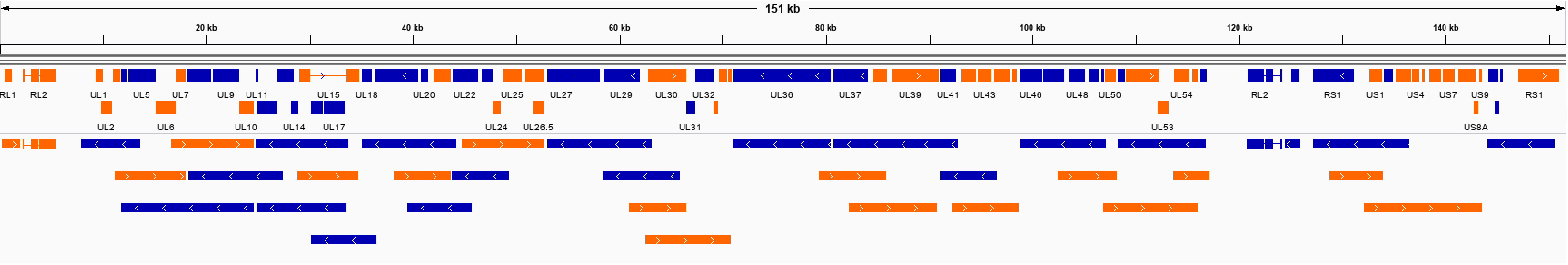

### Supplementary Figure 2

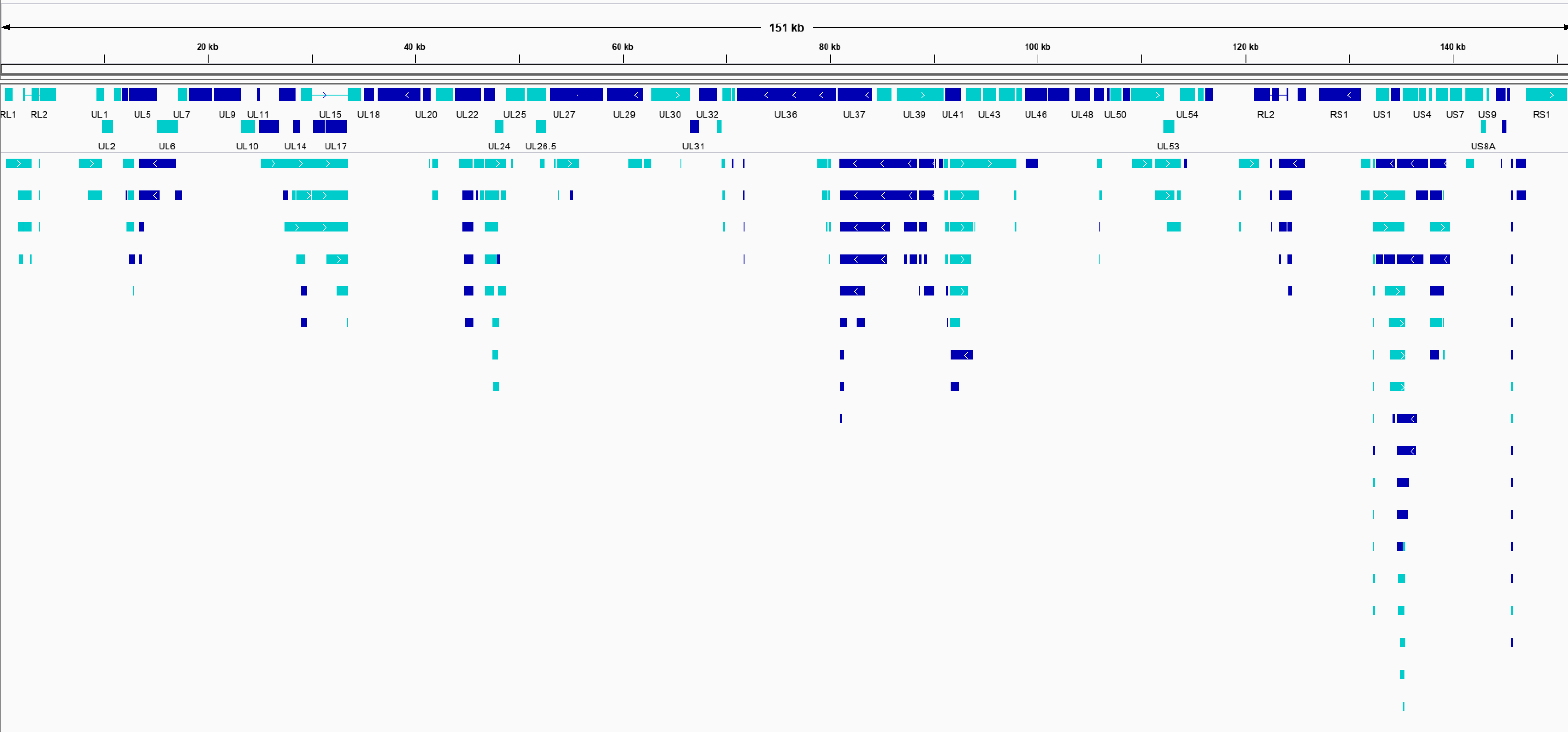
